# The GEN-ERA toolbox: unified and reproducible workflows for research in microbial genomics

**DOI:** 10.1101/2022.10.20.513017

**Authors:** Luc Cornet, Benoit Durieu, Frederik Baert, Elizabet D’hooge, David Colignon, Loic Meunier, Valérian Lupo, Ilse Cleenwerck, Heide-Marie Daniel, Leen Rigouts, Damien Sirjacobs, Stéphane Declerck, Peter Vandamme, Annick Wilmotte, Denis Baurain, Pierre Becker

## Abstract

**Background:** Microbial culture collections play a key role in taxonomy by studying the diversity of their accessions and providing well characterized strains to the scientific community for fundamental and applied research. These microbial resource centers thus need to implement new standards in species delineation, including whole-genome sequencing and phylogenomics. In this context, the genomic needs of the Belgian Coordinated Collections of Microorganisms (BCCM) were studied, resulting in the GEN-ERA toolbox. The latter is a unified cluster of bioinformatic workflows dedicated to both bacteria and small eukaryotes (i.e. yeasts).

**Findings:** This public toolbox allows researchers without a specific training in bioinformatics to perform robust phylogenetic analyses. Hence, it facilitates all steps from genome downloading and quality assessment, including genomic contamination estimation, to tree reconstruction. It also offers workflows for average nucleotide identity comparisons and metabolic modeling.

**Technical details:** Nextflow workflows are launched by a single command and are available on the GEN-ERA GitHub repository (https://github.com/Lcornet/GENERA). All the workflows are based on Singularity containers to increase reproducibility.

**Testing:** The toolbox was developed for a diversity of microorganisms, including bacteria and fungi. It was further tested on an empirical dataset of 18 (meta)genomes of early-branching Cyanobacteria, providing the most up-to-date phylogenomic analysis of the *Gloeobacterales* order, the first group to diverge in the evolutionary tree of Cyanobacteria.

**Conclusion:** The GEN-ERA toolbox can be used to infer completely reproducible comparative genomic and metabolic analyses on prokaryotes and small eukaryotes. Although designed for routine bioinformatics of culture collections, it can also be useful for other applications, as shown by our case study on *Gloeobacterales*.

## Background

Genomics has revolutionized a number of research fields, including microbial taxonomy. Nowadays, genomes are frequently used for species delineation; the average nucleotide identity (ANI) comparisons becoming the new gold standard for bacterial and yeast taxonomy, replacing DNA-DNA hybridization experiments [1–4]. The Genome Taxonomy Database (GTDB) project demonstrates the usefulness of this approach by providing a prokaryotic taxonomy completely based on genome sequences [5–6]. Complementary to ANI, phylogenomics is also increasingly used to guide the taxonomy of microorganisms, notably small eukaryotes [7–9]. Phylogenomic studies are based on the analysis of hundreds to thousands of genes at once, outperforming single-gene phylogenies in terms of resolution and accuracy [10–12].

Microbial culture collections are public biological resource centers that preserve and distribute microorganisms for many purposes, such as industrial applications, quality controls, teaching activities or scientific research at large. They also play an important role in taxonomy, either by investigating the phylogeny of their own strains or by distributing them to taxonomists for a symbolic fee [13–14]. To enforce a correct taxonomy for their diverse microbial materials, culture collections have to integrate modern genomic practices. This task is not trivial since genomics is a rapidly changing field and the bioinformatic pipelines are constantly evolving. For instance, the evaluation of genomic contamination has evolved a lot during the last three years, with 11 new algorithms published [15]. The production of genome assemblies can also require advanced metagenomic methods, depending on the axenic level of the cultures [16] [17].

In 2016, a survey designed to evaluate the bioinformatic reproducibility in Science reported that 70% of researchers failed to reproduce genomic research from other scientists and that 50% failed to reproduce their own research [18]. The main source of computational irreproducibility was due to variations between operating systems, and (lack of) availability of software and databases [19]. These limitations can be overcome by the use of Singularity containers that package softwares in a frozen computational environment [20]. Nextflow is a Singularity-aware workflow system that is well suited to address the challenge of reproducibility [19].

The availability of reproducible genomic tools for taxonomic studies is relevant for microbial collections. In this context, the needs of five collections belonging to the Belgian Coordinated Collections of Microorganisms (BCCM) were addressed in the framework of the Belgian Science Policy (BelSPO) GEN-ERA project (https://bccm.belspo.be/content/bccm-collections-genomic-era). The latter aimed to establish modern genomic practices for improving the taxonomy of various types of microorganisms: moulds, yeasts, cyanobacteria, mycobacteria, and endosymbiotic bacteria/fungi. We report here the implementation of 13 Nextflow workflows, supported by 14 Singularity containers, which cover the most common genomic applications related to microbial taxonomy, including metabolic modeling. To our knowledge, GEN-ERA is the first unified publicly available toolbox designed for genomic studies on bacteria and yeasts. It is also designed to be used by collection personnel without deep knowledge of bioinformatics.

## Findings

Here, we only give an overview of the GEN-ERA toolbox (**Figure 1**), while detailed descriptions are provided in the Methods section.

**Figure 1:**
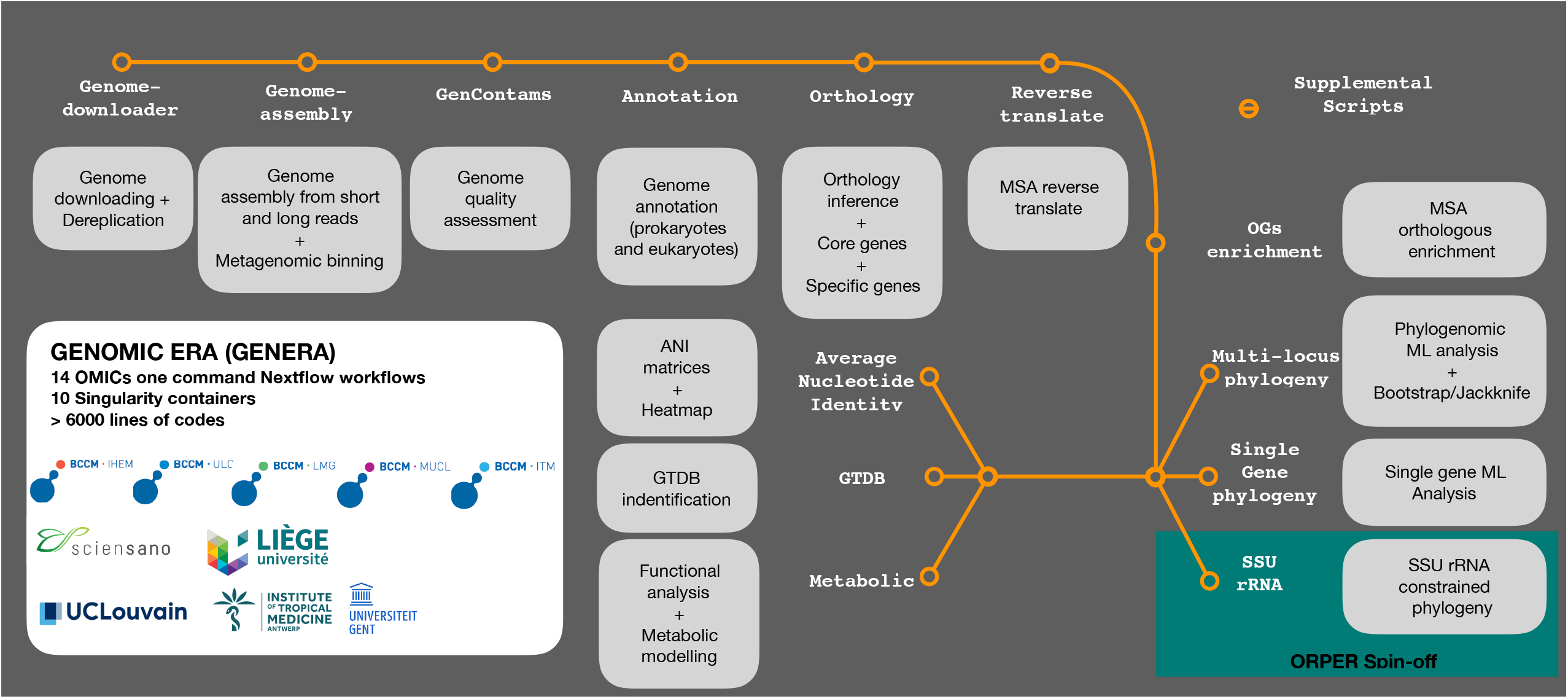
Overview of the GEN-ERA toolbox.

### GEN-ERA overview

#### Genome-related workflows

The first four workflows are related to genome acquisition and annotation. The first tool, ***Genome-downloader.nf***, automatically updates a local mirror of the NCBI Taxonomy [21] [22] at each run and then downloads the genomes according to this taxonomy. The user should specify the name of the group and the taxonomic rank (for instance, “Gloeobacterales” and “order”). The specification of the taxonomic rank makes ***Genome-downloader.nf*** resilient to future changes in the NCBI Taxonomy, as it happened recently [https://ncbiinsights.ncbi.nlm.nih.gov/2021/12/10/ncbi-taxonomy-prokaryote-phyla-added/].

The second tool, ***Assembly.nf***, is dedicated to genome production. This workflow can assemble genomes and metagenomes, not only from Illumina short reads but also PacBio or Nanopore long reads data, thanks to the use of SPAdes [23], metaSPAdes [24] and metaFlye [25]. An option for metagenomic binning, grouping contigs into individual metagenome-assembled genomes (MAGs), with MetaBAT2 [26] and CONCOCT [27], is provided too. These two binning algorithms are complementary, as CONCOCT is more efficient for eukaryotic data [28] while MetaBAT2 was pre-trained for prokaryotic sequences [26].

The third genome-related tool, ***GENcontams.nf***, is used for the estimation of genomic contamination and production of genome statistics. Contamination estimation (i.e., the inclusion of foreign DNA in a genome assembly) requires the use of multiple tools to recognise contaminants more accurately [15]. Indeed, some tools are dedicated to bacterial genomes (CheckM [29], GUNC [30]), others are specific to eukaryotes (EukCC [28]), and a few can work on both domains without the ability to perform interdomain detection (BUSCO [31]). In addition, Physeter [32] and Kraken2 [33] are two tools able to perform interdomain detection, allowing for instance the detection of eukaryotic contamination in bacteria (and vice versa). To facilitate the detection of contaminants, all these tools are implemented in ***GENcontams.nf***. Researchers interested in a better understanding of these tools can read the recent review on the detection of genomic contamination made by Cornet et al. [15]. Besides, the genome assembly quality assessment tool QUAST [34] is provided in ***GENcontams.nf*** for classical genome statistics.

The last tools of this section are related to genome annotation (i.e., protein prediction). The annotation of bacterial proteins is automatic in the different GEN-ERA workflows, but we nevertheless provide a Singularity container for bacterial protein prediction with Prodigal [35]. In opposition to bacteria, eukaryotic gene annotation is not automatic in the GEN-ERA suite, but two tools, ***AMAW*** [36] and ***BRAKER.nf***, are included for this usage. The workflow ***BRAKER.nf*** is able to download RNAseq evidence, based on a user-provided list, and to use proteins from OrthoDB [37] to annotate genomes with BRAKER2 [38]. In contrast, ***AMAW*** automatizes evidence collection based on the species name [36] and is dedicated to annotation of non-model organisms.

#### Phylogeny-related workflows

This section covers phylogenomic analysis from orthology inference to production of phylogenomic trees. The first workflow, ***Orthology.nf***, implements orthology inference. Bacterial genomes (or proteomes) and eukaryotic proteomes are the basis of ***Orthology.nf***. Two software tools can be used to compute orthologous groups (OGs) of proteins: OrthoMCL [39], available for prokaryotes only, and OrthoFinder [40], available for both domains. ***Orthology.nf*** automatically provides the core genes, shared by all the organisms in unicopy, and the specific genes, found only in a user-provided list of organisms. The OGs of proteins can be further enriched with orthologous sequences from new organisms by ***OGsEnrichment.nf***, using Forty-Two [41–42], available at https://metacpan.org/dist/Bio-MUST-Apps-FortyTwo). OGs can also be reverse translated by ***OGsRtranslate.nf***, using Leel ([43]; available at https://metacpan.org/dist/Bio-MUST-Apps-FortyTwo). Both protein and nucleotidic OGs can then be used for phylogenomic analysis with ***Phylogeny.nf***. This workflow implements phylogenomic inference using BMGE [44] for selection of unambiguously aligned sites, SCaFoS [45] for sequence concatenation, and RAxML [46] for tree reconstruction. With a user interface very similar to ***Phylogeny.nf***, both types of OGs can also be provided to ***PhylogenySingle.nf*** in order to compute single-gene trees with RAxML[46].

The last tool of this section is ***ORPER.nf***, which was published independently [47] and is designed to constrain an SSU rRNA phylogeny with a phylogenomic backbone [46]. This tool first produces a phylogenomic tree based on concatenated ribosomal proteins, extracted from public genomes, and then constraints the larger SSU rRNA phylogeny using this reference phylogenomic tree. This multi-locus constraint is used to reduce the inaccuracy of single-gene analyses [47]. ORPER permits to localize new lineages, based on SSU rRNA diversity, without sequenced genome or to identify genomes close to strains for which only SSU rRNA sequences are available.

#### Other workflows

Three additional workflows are provided in the GEN-ERA toolbox. The first one, ***ANI.nf***, computes average nucleotide distances between genomes using fastANI [48]. The second one, ***GTDB.nf***, uses GTDBTk [49] to classify prokaryotic genomes according to the Genome Taxonomy Database (GTDB) [5–6]. The last workflow, ***Metabolic.nf***, is dedicated to protein function annotation using Mantis [50], and metabolic modeling of prokaryotes using Anvi’o [51] with the Kyoto Encyclopedia of Genes and Genomes (KEGG) database as a reference [52].

#### Implementation

The workflows are developed with Nextflow workflow system [19] and are all supported by Singularity containers [20], except for the Mantis part of ***Metabolic.nf***, because it was technically not possible to include Mantis in a container. Instead, we documented how to use it from a conda environment. Each workflow is accompanied by a python script for parsing and formatting results, included in the containers. The workflows are provided to the users as programs and each includes a help section. They can be run with a single command, increasing the reproducibility of the analyses. The databases used by the different workflows (Table 1) are automatically downloaded at the first run of the workflow if not pre-installed by the user. The GEN-ERA toolbox (workflows, Singularity definition files, companion scripts) is freely available from the GitHub repository: https://github.com/Lcornet/GENERA. This repository includes a detailed user guide for each tool, focusing notably on HPC cluster usage.

**Table 1:** Purpose of the GEN-ERA tools along with their databases and availability of Singularity containers.

| **Tool** | **Purpose** | **Databases used** | **Availability of containers** |
| --- | --- | --- | --- |
| Genome-downloader.nf | Download of NCBI genomes and proteomes | NCBI Taxonomy<br>V:automatic setup | Yes |
| Assembly.nf | Assembly of (meta)genomes from short and long reads, binning of metagenomes | None | Yes |
| GENcontams.nf | Estimation of genome quality | NCBI Taxonomy VJune 13th 2021<br>GUNC: progenomes2.1<br>Physeter: Cornet et al., 2021<br>BUSCO db Vodb.10<br>Kraken db STD+<br>eukcc2_db_ver_1.1 | Yes |
| AMAW | Eukaryotic genome annotation | prot_dbEnsembl Protists, Fungi and Plants release 35.0 in combination with protist genomes available on the NCBI in March 2017<br>augustus_db VJune 28th 2021 | No |
| Braker.nf | Eukaryotic genome annotation | OrthoDB Vodb10<br>Augustusdb VJune 28th 2021 | No |
| Orthology.nf | Orthologous inference, delineation of core and specific genes. | NCBI Taxonomy VJune 13th 2021 | Yes |
| OGsEnrichment.nf | Orthologous enrichment of amino-acid OGs with sequences from genomes and proteomes. | NCBI Taxonomy<br>June 13th 2021 | Yes |
| OGsRtranslate.nf | Reverse translation of amino-acid OGs. | None | Yes |
| Phylogeny.nf | ML phylogenomic analysis, with bootstrap and jackknife replicates, of amino-acid and nucleotide sequences. | None | Yes |
| PhylogenySingle.nf | Single-gene ML phylogeny of amino-acid and nucleotide sequences. | None | Yes |
| ORPER.nf | SSU rRNA constrained ML phylogeny. | RiboDB | Yes |
| ANI.nf | Average nucleotide identity comparison. | None | Yes |
| GTDB.nf | Genome identification according to GTDB. | GTDB version Vr207 | Yes |
| Metabolic.nf | Functional and metabolic analyses. | MantisDB V1.5.4<br>KEGG version V202 | No |

#### Testing

The GEN-ERA toolbox was initially tested by the users from the BCCM involved in the GENERA project, who were thus considered as beta testers, on a SLURM-operated HPC system (durandal2/nic5, CÉCI-ULiège). These users were not advanced bioinformatics researchers and the user guide was developed based on their needs to ensure an easy-to-use toolbox. This toolbox was further tested on the *Gloebacterales* order (Cyanobacteria) as a case study. All command lines used for this test case are provided in Supplemental Note 1.

##### *Gloeobacterales* as a case study

Composed of thylakoid-less bacteria [53–54], *Gloeobacterales* are the most basal order of the Cyanobacteria phylum. Being the first group to have diverged, it is of particular interest for the study of cyanobacterial evolution. This order has long been represented by only two genomes (see for instance Cornet et al., 2018 [55] and Moore et al., 2019 [56] phylogenies). However, the diversity of the group was recently expanded with new genomes obtained from cultivated strains [57–58] and from metagenomes [53, 59–60]. *Gloeobacter* spp. strains were isolated from rock biofilms but the SSU sequences and metagenomes data show that they are widely distributed [53, 61]. For instance, the metagenomes of *Aurora vendensis* were isolated from the benthic microbial mats in an Antarctic lake [59] and the strain *Anthocerotibacter panamensis* from the surface-sterilized thallus of the hornwort *Leiosporoceros dussii* from Panama [58]. Here, we used the GEN-ERA toolbox to produce, in a completely reproducible manner, the most up-to-date phylogeny of the *Gloeobacterales* order, composed of eight (meta)genomes (Figure 2A, Supplemental File 1). In brief, we downloaded the genomes, estimated their contamination level, reassembled a genome deleted from the NCBI repository, then computed large amino acid and nucleotide phylogenomic analyses, both supported by bootstrap and jackknife resampling (Figure 2A, Supplemental File 1). Seven *Gloeobacterales* genomes were available on NCBI servers and were automatically downloaded by our tools (see Supplemental Note 1). One additional genome of *Gloeobacterales, Gloeobacteraceae* cyanobacterium ES-bin-313 from an Arctic Glacier [60], had been deleted from NCBI servers due to a low completeness. We re-assembled this genome from the raw reads and used the assembly in a phylogenomic analysis of the group for the first time. The automatization of the GEN-ERA workflows allowed us to automatically include all available strains in our phylogenies. The Supplementary figures S1-S4 showed two clusters, one with the (meta)genomes of *Gloeobacter* spp. and the other with the (meta)genomes of candidatus *A. vandensis* and *A. panamensis*, as expected [58]. We also used 566 SSU rRNA sequences from the SILVA repository [62] to estimate the sequencing level of the order by computing an SSU rRNA phylogeny constrained by the eight public genomes thanks to ORPER [47] (Figure 2B). The constrained SSU rRNA phylogeny revealed 11 sequences branching at a very basal position in the cyanobacterial tree, before any known *Gloeobacterales* genomes, an observation never made before, as far as we know. These sequences likely represent interesting targets for future whole genome sequencing projects. This confirms the interest of using ORPER to spot interesting SSU rRNA sequences, of which the organism would deserve a genome sequencing. We also applied ANI comparisons to the eight publicly available genomes and investigated the presence of biosynthesis KEGG pathways in *Gloeobacterales* and closely associated strains. Our results demonstrate the absence of one metabolic pathway in the *Gloeobacterales* order and of two pathways in the *Gloeobacter* group. The first pathway, absent in the whole *Gloeobacterales* order, is involved in the citrate cycle (Supplemental Note 1). Two other pathways involved in carotene and isoprenoid biosynthesis are absent from the *Gloeobacter* group but present in all other sampled Cyanobacteria, at the exception of the marine *Synechococcus sp*. PCC7336. (Figure 2C). *Anthocerotibacter panamensis* C109 is the only sampled cyanobacterium to present the archaeal (M00365) isoprenoid biosynthesis pathway (Figure 2C). This might result from a genuine lateral gene transfer, because the contamination level of this genome is very low (0.85 %). Detailed results and examples of the practical usage of the GEN-ERA toolbox are available in Supplemental File 1.

**Figure 2:**
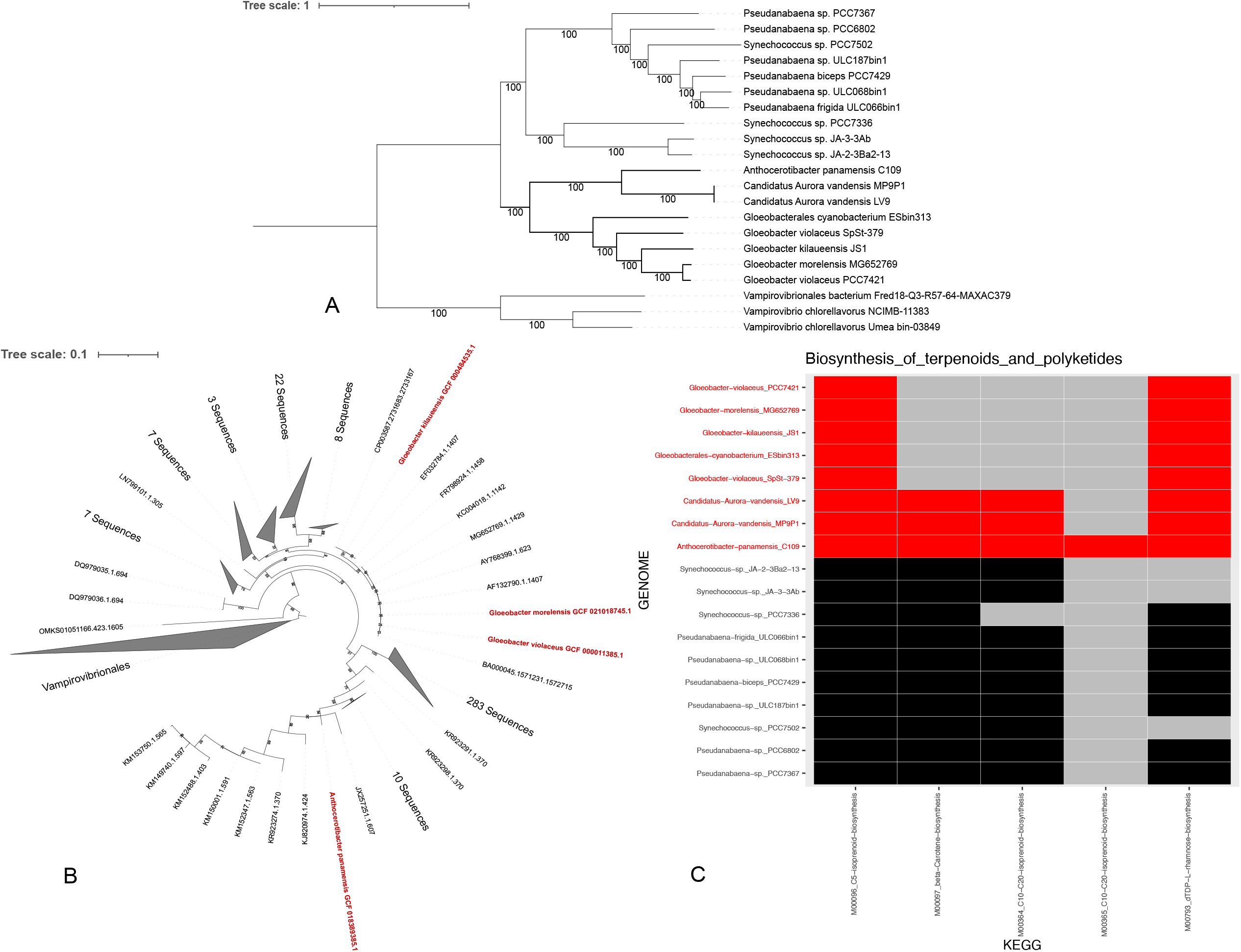
Results of the *Gloeobacterales* analysis. **A**. Phylogenomic analysis of the *Gloeobacterales* order, conducted on 198 core genes using DNA sequences. The tree was inferred with RAxML under the GTRGAMMA model on a supermatrix of 21 X 225,524 unambiguously aligned nucleotide positions. **B**. SSU rRNA phylogeny constrained by a phylogenomic analysis of ribosomal proteins, computed with ORPER. **C**. Metabolic modeling of *Gloeobacterales* and closely associated taxa. Detailed methods and results of the *Gloeobacterales* analysis are available in Supplemental File 1. *Gloeobacterales* are indicated in red.

## Methods

The versions of the programs used in the case study are provided below and correspond to the first public release of the GEN-ERA toolbox (Table 1).

### Genome-downloader.nf

A list of GCF accessions, from RefSeq [63–64], and GCA accessions, from GenBank [65–66] is created based on the assembly summary lists available on the NCBI FTP repository [22]. A local mirror of the NCBI Taxonomy is loaded with the script *setup-taxdir.pl* V0.212670 from the Bio-MUST-Core suite (available at https://metacpan.org/dist/Bio-MUST-Core). The taxonomic lineage, from phylum to species, of each genome is obtained based on the GCF/GCA number with the companion script *fetch-tax.pl* V0.212670 (also available at https://metacpan.org/dist/Bio-MUST-Core). Genomes are then downloaded according to the taxon name and taxonomic rank specified by the user. Priority is given to GCF over GCA assemblies for download. An optional dereplication of the genomes can be performed with *dRep* V3.0.0 [67] using the dereplicate option (with or without the ignoreGenomeQuality option). Finally, the proteins of the selected genomes can be downloaded if they exist on NCBI servers. https://github.com/Lcornet/GENERA/wiki/07.-Genome-downloader.

### Assembly.nf

This workflow can take as input both short (Illumina) and long reads (PacBio and Oxford Nanopore). Short reads are first trimmed and filtered to delete low-quality reads and adapters with *fastp* V0.23.1 [68], with default settings. If only short reads are provided, the assembly is performed with *SPAdes* V3.15.3 [23] with default settings. *metaSPAdes* V3.15.3 [24] is used if the metagenome option of the workflow is specified. If long reads are provided, the assembly can be done either with *Flye* V2.19.b1774 [25], with default settings, or *CANU* V2.3 [69], with the options stopOnLowCoverage=5 and cnsErrorRate=0.25. *Flye* V2.19.b1774 [25], with the meta option, is the only long-read assembler available with the metagenome option. An expected genome size should be provided by the user for all long-read assemblies. The polishing of such assemblies is carried out with *pilon* V1.24 [70], with default settings, after mapping of the short reads with *bwa mem* V0.7.17 [71] and *samtools* V1.13 [72]. The metagenomic binning to obtain individual Metagenome-Assembled Genomes (MAGs) is performed with MetaBAT2 V2.15.6 [26], with default settings, and/or *CONCOCT* V1.1 [27], with default settings too. The short-read coverage is provided as input for binning after mapping with *bwa mem* V0.7.17 [71] and *samtools* V1.13 [72]. Finally, a mapping of the contigs on a reference genome, not available for metagenomes, can be performed with RagTag V2.1.0 [73]. https://github.com/Lcornet/GENERA/wiki/08.-Genome-assembly.

### GENcontams.nf

This workflow estimates the level of genomic contamination with six different algorithms. The first tool is *CheckM* V1.1.3 [29], used with the lineage_wf option and the provided database. The second algorithm is *GUNC* V1.0.5 [30], with default settings, and is used with the database Progenomes 2.1 [74]. The third tool is *BUSCO* V5.3.0 [31], used in auto-lineage mode and with the provided database. The fourth tool is *Physeter* V0.213470 [32], a parser for *DIAMOND blastx* [75] reports. *Physeter* V0.213470 is used with the auto-detect option and with the database provided in Lupo et al. [32]. The fifth algorithm is *Kraken 2* V2.1.2 [33], used with default settings. The database of *Kraken 2* corresponds to the ‘PlusFP’ database downloaded from https://benlangmead.github.io/aws-indexes/k2. The sixth algorithm is EukCC [28], used with default settings and the provided database. Finally, statistics on the quality of genome assemblies are computed with *QUAST* V5.1.orc1 [34], with default settings. All the algorithms can be run independently but can also be used in one go to generate a summary table. The various databases of the different tools are automatically downloaded if not provided by the user. https://github.com/Lcornet/GENERA/wiki/09.-Genome-quality-assessment.

### BRAKER.nf

Eukaryotic genome annotation can be performed with *AMAW* [36], a MAKER2 [76] pipeline wrapper dedicated to non-model organisms and automating the orchestration of its internal annotation steps, as well as the collection of species-specific transcripts and phylogenetically related protein evidence data. *BRAKER 2* V2.1.6 [38] can also be used on eukaryotic genomes. Based on a user-provided list of RNASeq SRA numbers, the generation of transcript hints is performed by mapping the reads using *HISAT2* V9.2.1 [77] and *samtools* V1.13 [72], with default settings. Genomes of the OrthoDB [37] repository are used as protein evidence and are available in three different batches: fungi, protozoa and plants. https://github.com/Lcornet/GENERA/wiki/10.-Annotation.

### Orthology.nf

Orthology inference can be performed with OrthoFinder V2.5.4 [40], used with default settings, or with OrthoMCL [39] through the pangenomic pipeline of Anvi’o V7.1 [51]. The Anvi’o mode, available for prokaryotes only, requires the use of nine different scripts: anvi-script-reformat-fasta (with the options simplify-names and seq-type set to NT), anvi-gen-contigs-database (with default settings), anvi-run-ncbi-cogs (with default settings), anvi-gen-genomes-storage (with default settings), anvi-pan-genome (with the options mcl-inflation set to 10 and min-occurrence set to 2), anvi-get-sequences-for-gene-clusters (with default settings), anvi-script- add-default-collection (with default settings), anvi-summarize (with default settings) and anvi-compute-gene-cluster-homogeneity (with default settings). Orthology inference usually starts from complete proteomes. Nevertheless, prokaryotic genomes can be used, as prediction for prokaryotes with prodigal [35], is included in the workflow. In contrast, eukaryotic proteins should be provided by the user to Orthology.nf. After orthology inference, Orthology.nf can compute (optional) core genes. Core genes are considered here as unicopy genes shared by all organisms (and only these organisms) of a user-specified list, without exception. Another option allows the user to determine the specific genes, considered here as genes specific to a sub-list of organisms, without intruders. The main difference with core genes is that specific candidate OGs will undergo an orthologous enrichment by mining the genomes of all the organisms of the orthologous inference. This strategy is used in our analyses of the *Snodgrassella*-specific gene content [78] to prevent any orthologous delineation bias. Orthologous enrichment is performed with Forty-Two V0.212670 [41–42] (available at https://metacpan.org/dist/Bio-MUST-Apps-FortyTwo), with the same settings as ***OGsEnrichment.nf***. https://github.com/Lcornet/GENERA/wiki/11.-Orthology.

### OGsEnrichment.nf

This workflow can take as input amino acid OGs, as produced by ***Orthology.nf***. OGs can be aligned with *MUSCLE* V3.8.31 [79], with default values. The enriching sequences can come from genomes or proteomes. In both cases, BLAST banks are built with *makeblastdb* V2.10.0 [80]. The orthologous enrichment is performed with *Forty-Two* V0.212670 [41–42] (available at https://metacpan.org/dist/Bio-MUST-Apps-FortyTwo). *Forty-Two* V0.212670 is used with a BLAST e-value of 1e-05, a max_target_seqs of 10000, the templates_seg option set to no, the ref_org_mul set to 0.3, the ref_score_mul set to 0.99, the trim_homologues option set to on, the ali_keep_lengthened_seqs option set to keep and the ref_brh enabled. The default aligner is *BLAST* V2.10.0 but the user can also use *exonerate* V2.2.0. https://github.com/Lcornet/GENERA/wiki/13.-OGs-Enrichment.

### OGsRtranslate.nf

As for ***OGsEnrichment.nf***, OGs can be aligned with *MUSCLE* V3.8.31 [79], with default values. Protein sequence alignments are back-translated by capturing and aligning the corresponding DNA sequences with the program *Leel* V0.212670 [43] (available at https://metacpan.org/dist/Bio-MUST-Apps-FortyTwo). https://github.com/Lcornet/GENERA/wiki/12.-OGs-DNA-reverse-translate.

### Multi-locus Phylogeny.nf

This workflow takes as input OGs produced by ***Orthology.nf***, ***OGsEnrichment.nf*** or ***OGsRtranslate.nf***. The OGs can thus contain amino-acid or nucleotide sequences. As for the previous workflows, amino-acid OGs can be aligned with *MUSCLE* V3.8.31 [79], with default values. Nucleotide OGs are not aligned, as they are obtained by back-translating amino-acid alignments with ***OGsRtranslate.nf***. Unambiguously aligned positions in amino-acid OGs are selected with BMGE V1.12 [44], used with a “medium” mask, as specified in Bio-MUST-Core V0.212670 (available at https://metacpan.org/dist/Bio-MUST-Core). This selection is not performed on nucleotide OGs in order to preserve the codon phase. OGs are concatenated using *SCaFoS* V1.25 [45], with default settings. Finally, trees are inferred using *RAxML* V8.2.12 [46] with 100 bootstrap replicates under the PROTGAMMALGF model for proteins and the GTRGAMMA model for DNA sequences. DNA trees are computed either without a codon partition, or with a separate partition on the third codon position or based only on the two first positions. Beside these large phylogenomic analyses, the workflow also computes jackknife analyses. A hundred jackknife matrices are generated with the script *jack-ali-dir.pl* V0.212670 from Bio-MUST-Core (available at https://metacpan.org/dist/Bio-MUST-Core), using a width of 100 000 positions (modifiable by the user), and concatenated with *SCaFoS* V1.25 [45], as above. The trees are computed with *RAxML* V8.2.12 [46], as above (including codon partitions), but under the fast mode. The consensus trees, from the 100 trees obtained on the matrices, are produced with consense from the PHYLIP package V3.695 [81], used with default settings. https://github.com/Lcornet/GENERA/wiki/14.-Multi-locus-Maximum-Likelihood-Phylogeny. Two other workflows for phylogenetic analyses are available in the GEN-ERA toolbox: ***PhylogenySingle.nf*** and ***ORPER.nf***. ***PhylogenySingle.nf*** is a simpler version of ***Phylogeny.nf***, with the same alignment, filtering of unambiguous aligned positions and tree reconstruction settings, but for single-gene analyses. https://github.com/Lcornet/GENERA/wiki/15.-Single-locus-Maximum-Likelihood-Phylogeny. ***ORPER.nf***, designed for constrained SSU rRNA phylogenetic inference, has already been published separately [47].

### ANI.nf

***ANI.nf*** performs pairwise average nucleotide identity comparisons using *fastANI* V1.33 [48] in an all-versus-all mode, with default settings. A heatmap is then computed, according to a user-specified list of genomes, with *ggplot2* [82]. https://github.com/Lcornet/GENERA/wiki/17.-ANI.

### GTDB.nf

This workflow allows the identification of genomes according to the GTDB taxonomy [5–6]. ***GTDB.nf*** uses *GTDBTk* V2.2.0-r207 [49] using the classify_wf workflow, with default settings. https://github.com/Lcornet/GENERA/wiki/18.-GTDB.

### Metabolic.nf

***Metabolic.nf*** is the last workflow of the GEN-ERA toolbox. It has two modes: functional or modeling. The functional mode carries out a functional characterization of protein sequences using Mantis V1.5.4 [50], with default settings, whereas the modeling mode provides modeling of KEGG pathways [52], based on the presence of at least 60% of the genes involved in a pathway, for prokaryotic genomes. This mode uses the *anvi-estimate-metabolism* of Anvi’o V7.1 [51]. Presence/absence plots of KEGG pathways is then graphically represented with *ggplot2* [82], according to a user-specified list of genomes. https://github.com/Lcornet/GENERA/wiki/19.-Metabolic.

## Availability of supporting source code and requirements

- Project name: GEN-ERA
- Project home page: https://github.com/Lcornet/GENERA
- Operating system(s): Platform independent, Singularity containers
- Programming language: Nextflow and Python
- Other requirements: None

## Data Availability

The data used for *Gloeobacterales* analysis were downloaded from the NCBI SRA repository (SRR7539891, SRR12931219, SRR12931218).

## Declarations

## List of abbreviations

AA: Amino acid
ANI: Average Nucleotide Identity
BCCM: Belgian Coordinated Collections of Microorganisms
OGs: Orthologous Groups
GTDB: Genome Taxonomy Database
KEGG: Kyoto Encyclopedia of Genes and Genomes
MAGs: Metagenome-Assembled Genomes
ML: Maximum Likelihood
SSU rRNA: Small-subunit ribosomal RNA

## Ethics approval and consent to participate

Not applicable.

## Authors’ contributions

LC, DB, PB conceived the study. LC developed the Nextflow workflows and Singularity containers with the help of DC. LM developed AMAW. VL developed Physeter. LC, BD, FB, ED tested the workflows. LC ran *Gloeobacterales* analyses and drew the figures. LC, DB, PB wrote the manuscript with the help of DS, LR, IC, HMD, AW, SD, PV.

## Acknowledgements

We thank Olivier Mattelaer for his help with Singularity containers.

