## Supplementary material for "The GEN-ERA toolbox: unified and reproducible workflows for research in microbial genomics": Supp file 1

### Supplemental File 1

This Supplemental Note shows how to use the GEN-ERA toolbox on basal, early emerging cyanobacteria. This file provides command lines and examples of outputs for:

1. Genome download
2. Genome quality estimation
3. Genome assembly
    a. Short reads assembly
    b. Long reads assembly
    c. GTDB classification
4. Orthology inference
5. Genome annotation
6. Orthologous enrichment
7. Reverse translation
8. Phylogenetic inference
    a. Amino-acid phylogenomic analyses
    b. Nucleotide phylogenomic analyses
    c. Constrained SSU rRNA analyses
9. Average Nucleotide Identity comparisons
10. Metabolic modeling

#### 1. Genome download

The basal part of the cyanobacterial phylogeny is composed of organisms belonging to the *Gloeobacterales*, the group of interest. The first step was to download the genomes present on the NCBI servers [1] [2] for this group but also for other Cyanobacteria in the basal lineages of cyanobacterial radiation and genomes of *Vampirovibrionales* for the outgroup, as it has been done recently by Rahmatpour et al., 2021 [3] and by Sat et al., 2021 [4].

```
#Genome-downloading workflow
#PATH: Genome-DW/

#Get outgroup
#As no Vampirovibrionales are present in RefSeq,
since metagenomic samples are not allowed in RefSeq, only GenBank is used
$ nextflow run Genome-downloader.nf --taxolevel=order \
--group=Vampirovibrionales --refseq=no --genbank=yes \
--dRep=no --ignoreGenomeQuality=no --cpu=20;  \
mv Genome-downloader_output Genome-downloader_Vampirovibrionales

#Get the other Cyanobacteria
$ nextflow run Genome-downloader.nf --taxolevel=genus --group=Pseudanabaena
--genbank=yes --dRep=no --ignoreGenomeQuality=no --cpu=20; \
mv Genome-downloader_output Genome-downloader_Pseudanabaena
```

```
$ nextflow run Genome-downloader.nf --taxolevel=genus \
--group=Synechococcus --genbank=yes --dRep=no \
--ignoreGenomeQuality=no --cpu=20;  \
mv Genome-downloader_output Genome-downloader_Synechcoccus

#Get the Gloeobacterales
$ nextflow run Genome-downloader.nf --taxolevel=order \
--group=Gloeobacterales --genbank=yes --dRep=no \
--ignoreGenomeQuality=no --cpu=20;  \
mv Genome-downloader_output Genome-downloader_Gloeobacterales

#CPU time: 36m 30s
```

These commands downloaded 19 genomes of *Vampirovibrionales*, 58 genomes of *Pseudanabaena*, 414 genomes of *Synechococcus* and 7 genomes of *Gloeobacterales*.

We selected the whole *Gloeobacterales* group as well as other basal Cyanobacteria representatives based on the phylogeny built by Cornet et al., 2018 [5]. This filtration of genomes can be done using the file "**Genomes.taxonomy",** by giving the lineage for each downloaded genome file present in each folder.

```
#Useful Synechococcus genomes
GCF_000013205.1 Synechococcus sp._GCF_000013205.1       \
GCF_000013205.1 Cyanobacteria; undef; Synechococcales; Synechococcaceae;
Synechococcus; Synechococcus sp. JA-3-3Ab
GCF_000013225.1 Synechococcus sp._GCF_000013225.1       \
GCF_000013225.1 Cyanobacteria; undef; Synechococcales; Synechococcaceae;
Synechococcus; Synechococcus sp. JA-2-3B'a(2-13)
GCF_000332275.1 Synechococcus sp._GCF_000332275.1       \
GCF_000332275.1 Cyanobacteria; undef; Synechococcales; Synechococcaceae;
Synechococcus; Synechococcus sp. PCC 7336
GCF_000317085.1 Synechococcus sp._GCF_000317085.1       \
GCF_000317085.1 Cyanobacteria; undef; Synechococcales; Synechococcaceae;
Synechococcus; Synechococcus sp. PCC 7502

#Useful Pseudanabaena genomes
GCA_003249035.1 Pseudanabaena sp._GCA_003249035.1   \
GCA_003249035.1 Cyanobacteria; undef; Pseudanabaenales; Pseudanabaenaceae;
Pseudanabaena; Pseudanabaena sp.
GCA_003242085.1 Pseudanabaena frigida_GCA_003242085.1  \
GCA_003242085.1 Cyanobacteria; undef; Pseudanabaenales; Pseudanabaenaceae;
Pseudanabaena; Pseudanabaena frigida
GCA_003249015.1 Pseudanabaena sp._GCA_003249015.1 \
GCA_003249015.1 Cyanobacteria; undef; Pseudanabaenales; Pseudanabaenaceae;
Pseudanabaena; Pseudanabaena sp.
GCF_000332215.1 Pseudanabaena biceps_GCF_000332215.1 \
GCF_000332215.1 Cyanobacteria; undef; Pseudanabaenales; Pseudanabaenaceae;
Pseudanabaena; Pseudanabaena biceps
```

```
GCF_000317065.1 Pseudanabaena sp._GCF_000317065.1 \
GCF_000317065.1 Cyanobacteria; undef; Pseudanabaenales; Pseudanabaenaceae;
Pseudanabaena; Pseudanabaena sp. PCC 7367
GCF_000332175.1 Pseudanabaena sp._GCF_000332175.1 \
GCF_000332175.1 Cyanobacteria; undef; Pseudanabaenales; Pseudanabaenaceae;
Pseudanabaena; Pseudanabaena sp. PCC 6802
```

Note:
In the GEN-ERA tools, bacterial genomes can be used for phylogeny. This is not the case for eukaryotes, where proteins have to be provided. It is possible to download the proteins of the genomes with the option --prot=yes of ***Genome-downloader.nf***.

#### 2. Genome quality estimation

The genomic contamination in the 89 downloaded genomes was estimated by CheckM [6] and GUNC [7]. CheckM can detect redundant contaminations, while GUNC is genome-wide and can also detect non-redundant contamination [6] [7] [8]. It is thus useful to use both algorithms. Eukaryotic contamination, which cannot be detected by these packages, is estimated by Physeter [9] and Kraken2 [10].

```
#GENcontams workflow
#PATH:  Quality/

#CheckM
$ nextflow run GENcontams.nf --genomes=GENERA-input --mode=checkm \
--ext=fna --cpu=20 --dbdir=/data/GENERA/CONTAMS  \
-- taxdump=/data/GENERA/CONTAMS/taxdump/ \
--taxlevel=phylum; mv GENERA-contams GENERA-contams_checkm

#GUNC
$ nextflow run GENcontams.nf --genomes=GENERA-input \
--mode=gunc --ext=fna --cpu=20 \
--dbdir=/data/GENERA/CONTAMS
--taxdump=/data/GENERA/CONTAMS/taxdump/ --taxlevel=phylum; \
mv GENERA-contams GENERA-contams_gunc

#KRAKEN 2
$ nextflow run GENcontams.nf --genomes=GENERA-input --mode=kraken \
--ext=fna --cpu=20 \
--dbdir=/data/GENERA/CONTAMS \
--taxdump=/data/GENERA/CONTAMS/taxdump/ \
--taxlevel=phylum; mv GENERA-contams GENERA-contams_kraken

#Physeter
$ nextflow run GENcontams.nf --genomes=GENERA-input --mode=physeter \
--ext=fna --cpu=20  --dbdir=/data/GENERA/CONTAMS \
--taxdump=/data/GENERA/CONTAMS/taxdump/ --taxlevel=phylum; \
mv GENERA-contams GENERA-contams_physeter
```

**#CPU time: 7h 55m 08s**

No evidence of eukaryotic contaminants was detected by Physeter and Kraken.
The results of CheckM and GUNC are shown in Table S1.
Genomes above contamination (> 5%) or below completeness (< 90%) thresholds, as defined by Bowers et al., 2017 [11], are indicated by a star (*).

| Genome | CK compl | CK_ contam | CSS | RRS | GUNC contam |
|---|---|---|---|---|---|
| Anthocerotibacter panamensis_GCF_018389385.1 | 98.29 | 0.85 | 0 | 0 | True |
| Aurora vandensis_GCA_013285555.1 | 93.16 | 1.71 | 0.09 | 0.08 | 0.46 |
| Gloeobacter kilaueensis_GCF_000484535.1 | 98.29 | 1.14 | 0.0 | 0.0 | 0.98 |
| Gloeobacter morelensis_GCF_021018745.1 | 99.15 | 0.85 | 0.06 | 0.07 | 0.79 |
| Gloeobacter violaceus_GCF_000011385.1 | 99.15 | 0.85 | 0.0 | 0.0 | 0.96 |
| Gloeobacterales cyanobacterium_GCA_014379585.1 | 95.73 | 4.56 | 0.2 | 0.04 | 0.39 |
| *Gloeobacterales cyanobacterium_GCA_014380935.1 | 92.32 | 6.50 | 0.19 | 0.05 | 0.55 |
| Pseudanabaena biceps_GCF_000332215.1 | 99.29 | 1.18 | 0.18 | 0.25 | 0.48 |
| Pseudanabaena frigida_GCA_003242085.1 | 98.82 | 0.47 | 0.09 | 0.28 | 0.5 |
| Pseudanabaena sp._GCA_003249015.1 | 99.29 | 0.47 | 0.07 | 0.25 | 0.72 |
| Pseudanabaena sp._GCA_003249035.1 | 97.09 | 0.71 | 0.07 | 0.2 | 0.51 |
| Pseudanabaena sp._GCF_000317065.1 | 98.23 | 0.59 | 0.0 | 0.0 | 0.93 |
| Pseudanabaena sp._GCF_000332175.1 | 99.76 | 0.71 | 0.0 | 0.0 | 0.98 |
| Synechococcus sp._GCF_000013205.1 | 100.00 | 1.32 | 0.0 | 0.0 | 0.96 |
| Synechococcus sp._GCF_000013225.1 | 100.00 | 0.00 | 0.0 | 0.0 | 0.96 |
| Synechococcus sp._GCF_000317085.1 | 99.76 | 0.00 | 0.0 | 0.0 | 0.95 |
| Synechococcus sp._GCF_000332275.1 | 100.00 | 4.39 | 0.0 | 0.0 | 0.96 |
| uncultured Vampirovibrio_GCA_910577395.1 | 91.45 | 1.42 | 0.01 | 0.57 | 0.44 |
| uncultured Vampirovibrio_GCA_910577515.1 | 92.31 | 1.14 | 0.0 | 0.0 | 0.46 |
| uncultured Vampirovibrio_GCA_910577635.1 | 90.31 | 0.00 | 0.0 | 0.0 | 0.44 |
| uncultured Vampirovibrio_GCA_910578465.1 | 92.31 | 1.14 | 0.0 | 0.0 | 0.46 |
| uncultured Vampirovibrio_GCA_910579975.1 | 90.60 | 0.09 | 0.12 | 0.2 | 0.62 |
| uncultured Vampirovibrio_GCA_910583725.1 | 90.60 | 1.14 | 0.01 | 0.58 | 0.42 |
| uncultured Vampirovibrio_GCA_910583875.1 | 91.45 | 1.99 | 0.01 | 0.47 | 0.46 |

| | | | | | |
|---|---|---|---|---|---|
| uncultured Vampirovibrio_GCA_910584405.1 | 90.60 | 0.85 | 0.21 | 0.39 | 0.6 |
| ->Vampirovibrio chlorellavorus_GCA_001858525.1 | 95.73 | 1.71 | 0.07 | 0.53 | 0.27 |
| *Vampirovibrio chlorellavorus_GCA_003149345.1 | 95.73 | 5.37 | 0.06 | 0.53 | 0.26 |
| Vampirovibrio chlorellavorus_GCA_003149375.1 | 94.87 | 1.71 | 0.02 | 0.57 | 0.31 |
| Vampirovibrio chlorellavorus_GCA_903826385.1 | 95.73 | 0.85 | 0.01 | 0.57 | 0.3 |
| ->Vampirovibrio chlorellavorus_GCA_903835245.1 | 95.73 | 0.85 | 0.01 | 0.57 | 0.3 |
| Vampirovibrio chlorellavorus_GCA_903839075.1 | 95.73 | 0.85 | 0.01 | 0.57 | 0.3 |
| Vampirovibrio chlorellavorus_GCA_903866275.1 | 95.73 | 0.85 | 0.01 | 0.57 | 0.3 |
| *Vampirovibrio sp._GCA_019634025.1 | 80.34 | 1.71 | 0.08 | 0.58 | 0.28 |
| Vampirovibrio sp._GCA_021323475.1 | 92.31 | 1.71 | 0.09 | 0.6 | 0.28 |
| ->Vampirovibrionales bacterium_GCA_016712355.1 | 95.73 | 0.85 | 0.08 | 0.61 | 0.24 |
| *Vampirovibrionales bacterium_GCA_021299615.1 | 75.77 | 0.85 | 0.02 | 0.63 | 0.28 |

**Table S1: Results of contamination estimation of NCBI genomes by CheckM (CK) and GUNC. CCS is the GUNC clade_separation_score. RRS is the GUNC reference_representation_score.**

The 4 genomes of low quality are discarded from further analyses. We only retain the 3 best *Vampirovibrionales* as the outgroup (indicated by an arrow (->) in Table S1).

### 3. Genome assembly

Aside from the genomes available on the NCBI servers, the genome of *Gloeobacter violaceus SpSt-379* (GCA_011331945.1) has been deleted from GenBank. Nevertheless, the raw reads of this genome are still available on the NCBI SRA servers (SRR7539891).
We assembled this genome in metagenomic mode and produced MAGs by binning as the BioProject mentions metagenomes in the sequencing section. In order to demonstrate the assembly of long reads using the GEN-ERA toolbox, we also reassembled *Gloeobacter morelensis_GCF_021018745.1.* This genome is the best long-read assembly, according to contamination values, among the *Gloeobacterales* (Illumina reads: SRR12931219, Nanopore reads: SRR12931218).

```
#Assembly
#PATH:  Assembly/

#Assembly of Gloeobacter violaceus SpSt-379
$ nextflow run Assembly.nf --shortreadsR1=SRR7539891_1.fastq \
--shortreadsR2=SRR7539891_2.fastq --metagenome=yes --binner=all --cpu=20
```

```
$ mv GENERA-assembly GENERA-assembly_St-379
#Production of 1 Metabat bin and 35 CONCOCT bins
#CPU time: 2m 38s

#Re-assembly of Gloeobacter morelensis
$ nextflow run Assembly.nf --shortreadsR1=SRR12931219_1.fastq \
--shortreadsR2=SRR12931219_2.fastq \
--ontreads=SRR12931218_1.fastq \
--genomeSIZE=5m --metagenome=no --cpu=20
$ mv GENERA-assembly GENERA-assembly_more
#Production of one genome
#CPU-time: 19m 54s
```

#### 3.1 GTDB classification

GTDBtk was used on the 35 CONCOCT bins to identify the *Gloeobacter* bin.

```
#GTDB on CONCOCT bins
#PATH:
$ /Supplemental-scripts/change-ext.py --currentext=fasta --newext=fn
$ nextflow run GTDB.nf --genome=genome --cpu=20
#CPU time: 4m 45s
```

Only one bin was affiliated to *Gloeobacter.*

```
#Result
CONCOCT_bin-1
d__Bacteria;p__Cyanobacteria;c__Cyanobacteriia;o__Gloeobacterales;\
f__Gloeobacteraceae;g__;s__
```

#### 3.2 Quality estimation of the assembled genomes

In this section, we estimated the contamination of the three newly assembled genomes, namely 1 MetaBAT2 bin, 1 CONCOCT bin for *Gloeobacter violaceus SpSt-379* and 1 genome for *Gloeobacter morelensis*.

```
#Quality estimation with CheckM and GUNC
#PATH:  Assembly/Quality/
$ nextflow run GENcontams.nf --genomes=GENERA-input \
--mode=checkm --ext=fasta --cpu=20 --dbdir=/data/GENERA/CONTAMS \
--taxdump=/data/GENERA/CONTAMS/taxdump/ --taxlevel=phylum
$ mv GENERA-contams GENERA-contams_checkm
$ nextflow run GENcontams.nf --genomes=GENERA-input --mode=gunc \
--ext=fasta --cpu=20 --dbdir=/data/GENERA/CONTAMS \
--taxdump=/data/GENERA/CONTAMS/taxdump/ --taxlevel=phylum
$ mv GENERA-contams GENERA-contams_gunc
#CPU time: 4m 42s
```

The results indicate an assembly quality of *Gloeobacter morelensis* similar to GCF_021018745.1, with the same values of CheckM completeness and contamination. As this new genome is nearly the same as the NCBI genome, we will retain GCF_021018745.1 in the subsequent analyses. The workflow gives an assembly for *Gloeobacter violaceus* SpSt-379 with a low completeness (<78%) but with few contamination.

| Genome | CK compl | CK_ contam | CSS | RRS | GUNC contam |
|---|---|---|---|---|---|
| CONCOCT_bin-1 | 99.15 | 0.85 | 0.11 | 0.03 | 0.75 |
| METABAT_bin-1 | 64/16 | 1.14 | 0.23 | 0.05 | 0.42 |

**Table S2: Results of contamination estimation after re-assembly by CheckM (CK) and GUNC. CCS is the GUNC clade_separation_score. RRS is the GUNC reference_representation_score.**

#### 4. Orthology inference

We voluntarily did not include the new assembly obtained above to demonstrate the orthologous enrichment later in the Supplemental Note. We ran the ***Orthology.nf*** workflow on 20 public genomes. We asked for the inference of core genes among orthologous groups (OGs).

```
#Orthology workflow
#PATH:  Orthology/
$ nextflow run Orthology.nf --infiles=infiles --mode=inference --core=yes \
--corelist=corelist --specific=no --anvio=yes --type=nucleotide --cpu=20
#CPU time: 2h 19m 54s
#This analysis produced only 16 core genes
#The outgroup Vampirovibrionales might be a little bit too distant to
catch enough gene with the default value
#We ran the pipeline again, without adding Vampirovibrionales in the core list,
but allowing them to be part of genes, not mandatory
#We also relaxed the anvio index (from 0.8 to 0.7) to allow more variation
between sequences in the core genes
$ mv corelist corelist-full #Backup the first core list with the 20 genomes
$ wc -l corelist #The new core list contains 17 genomes, without the three outgoup
#We used the REDO folder present in the output of Orthology.nf to run
only the core part of the workflow, with OG mode
$ mv infiles infiles-genomes
$ mv GENERA-Orthology/REDO/* infiles/
$ nextflow run Orthology.nf --infiles=infiles --mode=OG --core=yes \
--corelist=corelist --coreunwanted=3 \
--coreminfunctionalindex=0.7 --coremingeometricindex=0.7 --specific=no \
```

```
--anvio=yes --type=nucleotide --cpu=20
#CPU time: 19m 23s
#252 core genes obtained
```

252 core genes, present in all oxygenic Cyanobacteria in unicopy, are obtained with this workflow and will be used later for phylogeny.

Note:
The workflow is run with Anvi'o [12] in order to benefit from the structural and geometric metrics. For eukaryotes, as Anvi'o is not available, OrthoFinder [13] should be used instead.

#### 4. Genome annotation

We used Prodigal for protein prediction of the genomes.

```
#Annotation
#PATH: Annotation/
$ find *.fna > list
$ sed -i -e 's/.fna//g' list
$ for f in `cat list`; do singularity exec --bind
/scratch/ulg/bioec/lcornet/GLOEO/Annotation:/mnt \
/scratch/ulg/GENERA/prodigal-2.6.3.sif prodigal -i /mnt/$f.fna \
-o /mnt/GENERA.out -a /mnt/$f.faa -d /mnt/$f.genes.fna; done
#CPU time: 2m 03s
```

#### 5. Orthologous enrichment

We used the proteomes obtained from the Annotation section to perform orthologous enrichment of OGs with the sequences from the new assembly. The proteomes of the NCBI genomes were used as representative organisms. The proteome of *Gloeobacter violaceus SpSt-379* (CONCOCT-bin1) was used for enrichment.

```
#OGsEnrichment
#PATH: /scratch/ulg/bioec/lcornet/GLOEO/OGsEnrichment
$ nextflow run OGsEnrichment.nf --OG=OGs --representative=representative \
--org=org --ftaligner=blast --align=no --cpu=20
#CPU time: 26m 14s
```

218 out of the 252 core genes were enriched with sequences from *Gloeobacter violaceus* SpSt-379. 54 out of these 218 OGs have more than one sequence for *Gloeobacter violaceus SpSt-379* and will not be used in the analysis. A total of 198 OGs are thus available (252 minus 54) for phylogenomic inference.

Note: The OGs were not aligned before the enrichment because the alignment was done by Anvi'o. If OrthoFinder is used in **Orthology.nf**, the alignment option should be activated.

### 6. Reverse translation

The enriched OGs were reverse translated back to DNA to be used for phylogenetic inference.

```
#Rtranslate
#PATH: Rtranslate/
$nextflow run  OGsRtranslate.nf --OG=OGs --bank=bank  --ext=faa --align=no
#CPU time: 37m 55s
```

### 7. Phylogenetic inference

#### 7.1 Amino-acid phylogenomic inference

The amino-acid phylogenomic tree was computed on the 198 enriched protein OGs.

```
#Phylogenomic AA
#PATH: /scratch/ulg/bioec/lcornet/GLOEO/Phylo-AA
$ nextflow run Phylogeny.nf --OG=OGs --IDM=file.idm --cpu=20 \
--jackk=yes --align=no --width=50000 --mode=prot
#CPU time: 2d 4h 17m 23s
```

Two trees (one bootstrap and one jackknife) were produced.

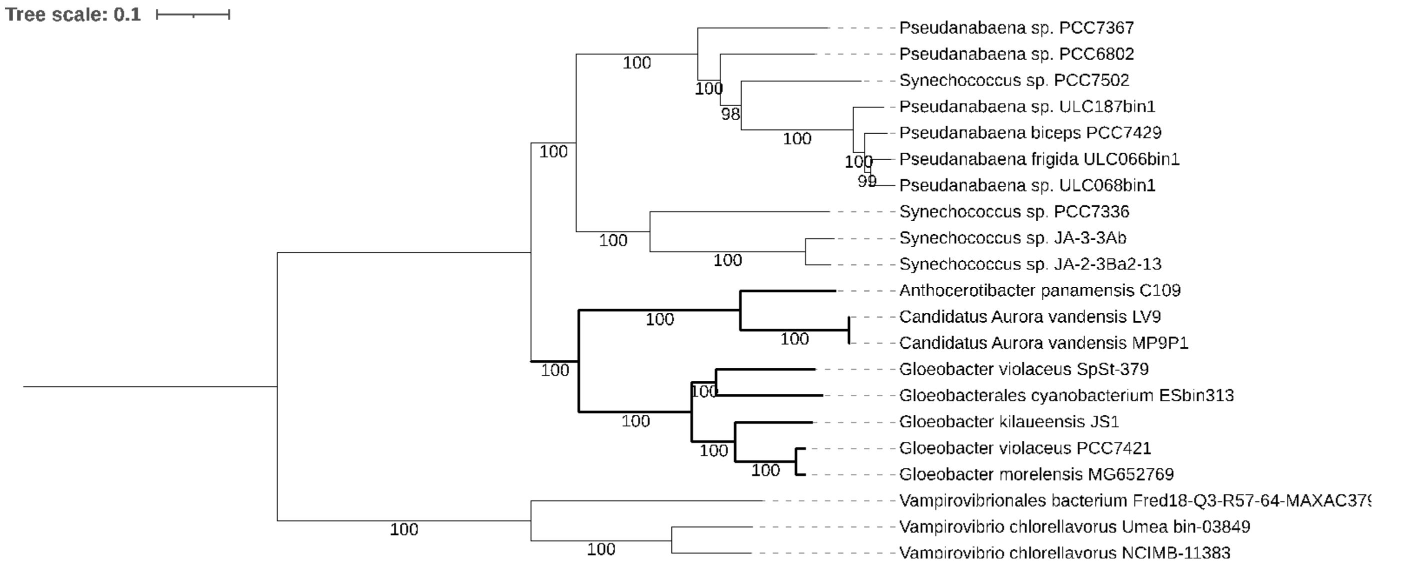


**Figure S1: Amino-acid phylogenomic analysis conducted on 198 genes. The tree was inferred with RAxML [14] under the PROTGAMMALGF model with 100 bootstrap replicates on a supermatrix of 21 X 65,143 unambiguously aligned positions. The *Gloeobacterales* group is indicated with bold branches.**

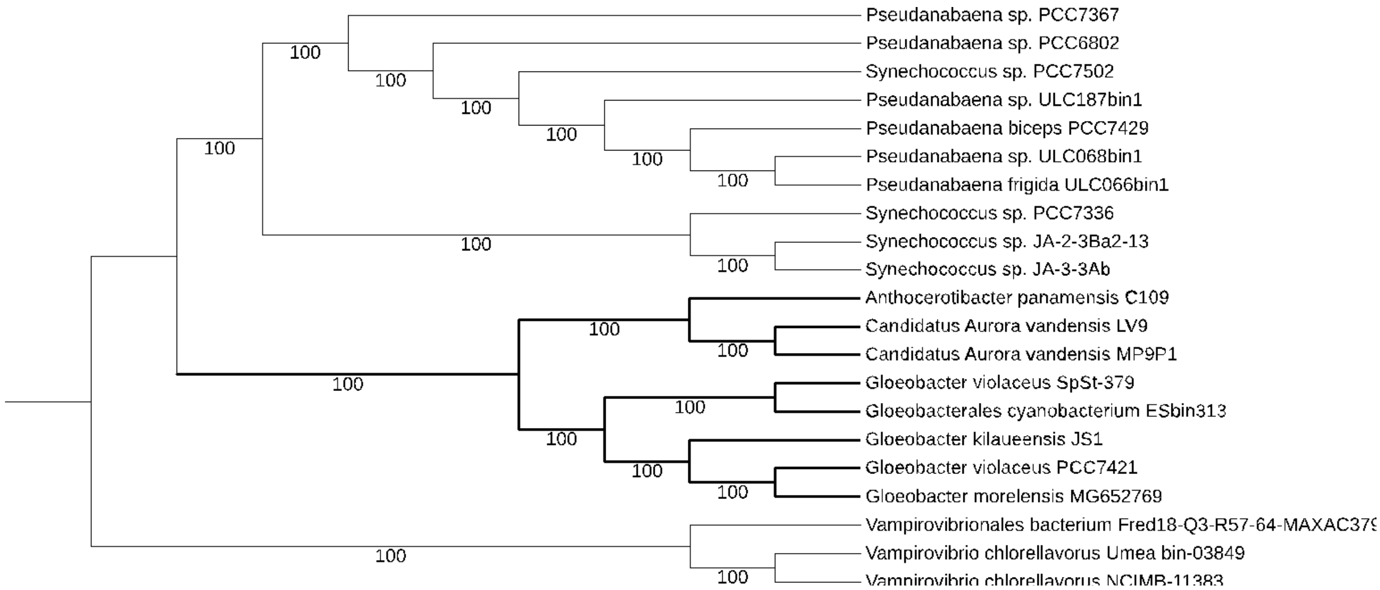


**Figure S2: Amino-acid phylogenomic analysis conducted on 198 genes. The tree was inferred with RAxML [14] under the PROTGAMMALGF using 100 x 50,000-AA random concatenations of genes produced by SCaFoS [15]. The *Gloeobacterales* group is indicated with bold branches.**

The two trees, bootstrap and jackknife, show the same phylogeny of the *Gloeobacterales*, both with maximum support.

#### 7.1 Nucleotide phylogenomic inference

The nucleotide phylogenomic analysis was computed from the 198 enriched DNA OGs.

```
#Phylogenomic NT
#PATH: /scratch/ulg/bioec/lcornet/GLOEO/Phylo-NT
$ nextflow run Phylogeny.nf --OG=OGs --IDM=file.idm --cpu=20 \
--jackk=yes --align=no --width=80000 --mode=DNA --ext=fna
#CPU time: 10h 59m 36s
```

Six trees were produced, in bootstrap and in jackknife: trees without codon partition, trees with a partition on codon position 3, trees based on only codon positions 1&2. We will only show here the trees based on codon positions 1&2.

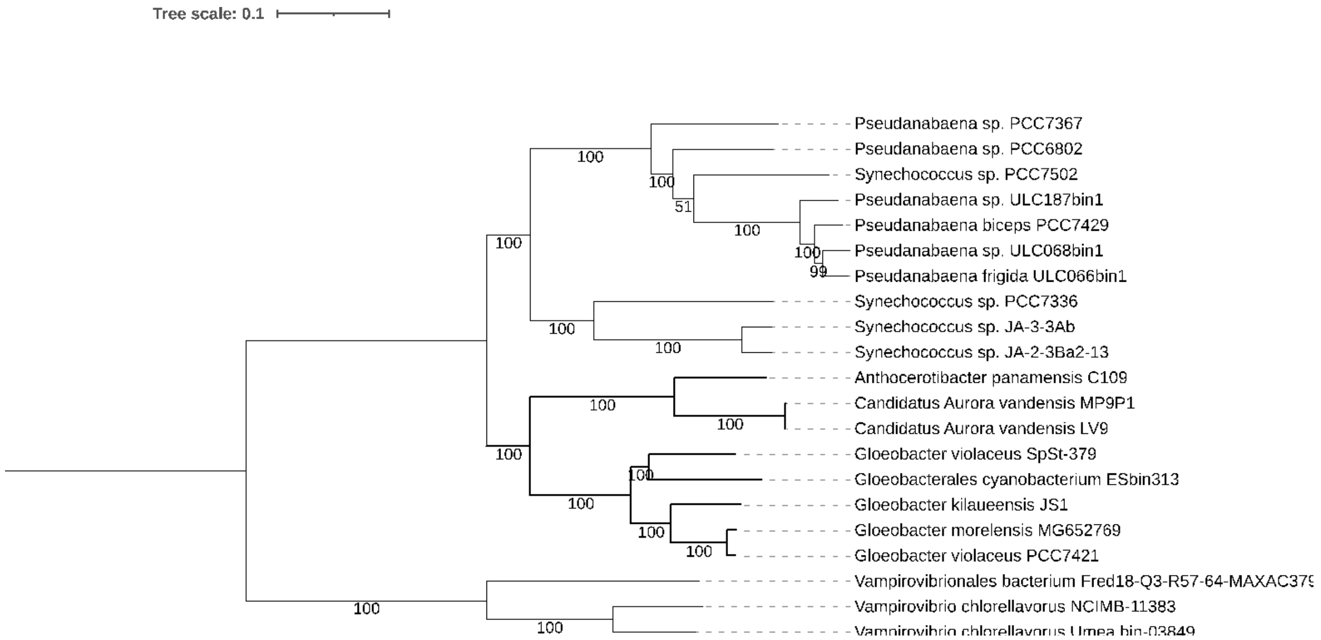


**Figure S3: Nucleotide phylogenomic analysis conducted on 198 genes. The tree was inferred with RAxML [14] under the GTRGAMMA model with 100 bootstrap replicates on a supermatrix of 21 X 225,524 positions. The *Gloeobacterales* group is indicated with bold branches.**

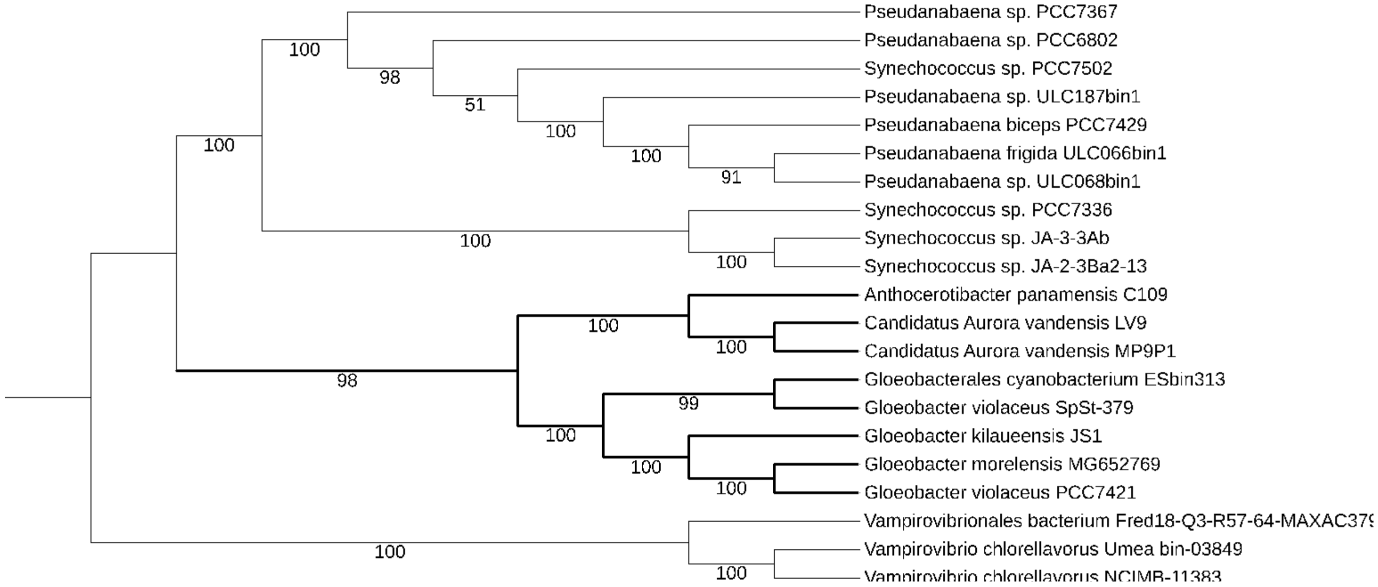


**Figure S4: Nucleotide phylogenomic analysis conducted on 198 genes. The tree was inferred with RAxML [14] under the GTRGAMMA model using 100 x 80,000 random concatenations of genes produced by SCaFoS [15]. The *Gloeobacterales* group is indicated with bold branches.**

The two trees show the same topology of the *Gloeobacterales* group, with support above 98 for all nodes.

#### 7.1 SSU rRNA constrained phylogenomic inference

566 SSU rRNA sequences affiliated to the *Gloeobacterales* order were downloaded from the SILVA servers (https://www.arb-silva.de) [16] on June 17th, 2022. We used ORPER [17] to compute an SSU rRNA phylogeny constrained by a phylogenomic analysis based on ribosomal proteins. ORPER permits researchers to quickly get an overview of the sequencing coverage of the order while reducing the uncertainty of SSU rRNA phylogeny [17].

```
#Constrained phylogeny
#PATH: /scratch/ulg/bioec/lcornet/GLOEO/ORPER
$ nextflow run ORPER.nf --reftaxolevel=order --refgroup=Gloeobacterales \
--refgenbank=yes --outgroup=Vampirovibrionales --outtaxolevel=order \
--outgenbank=yes --cpu=20 --SSU=silva.fasta --cdhit=yes --drep=no
#CPU time: 1h 12m 48s
```

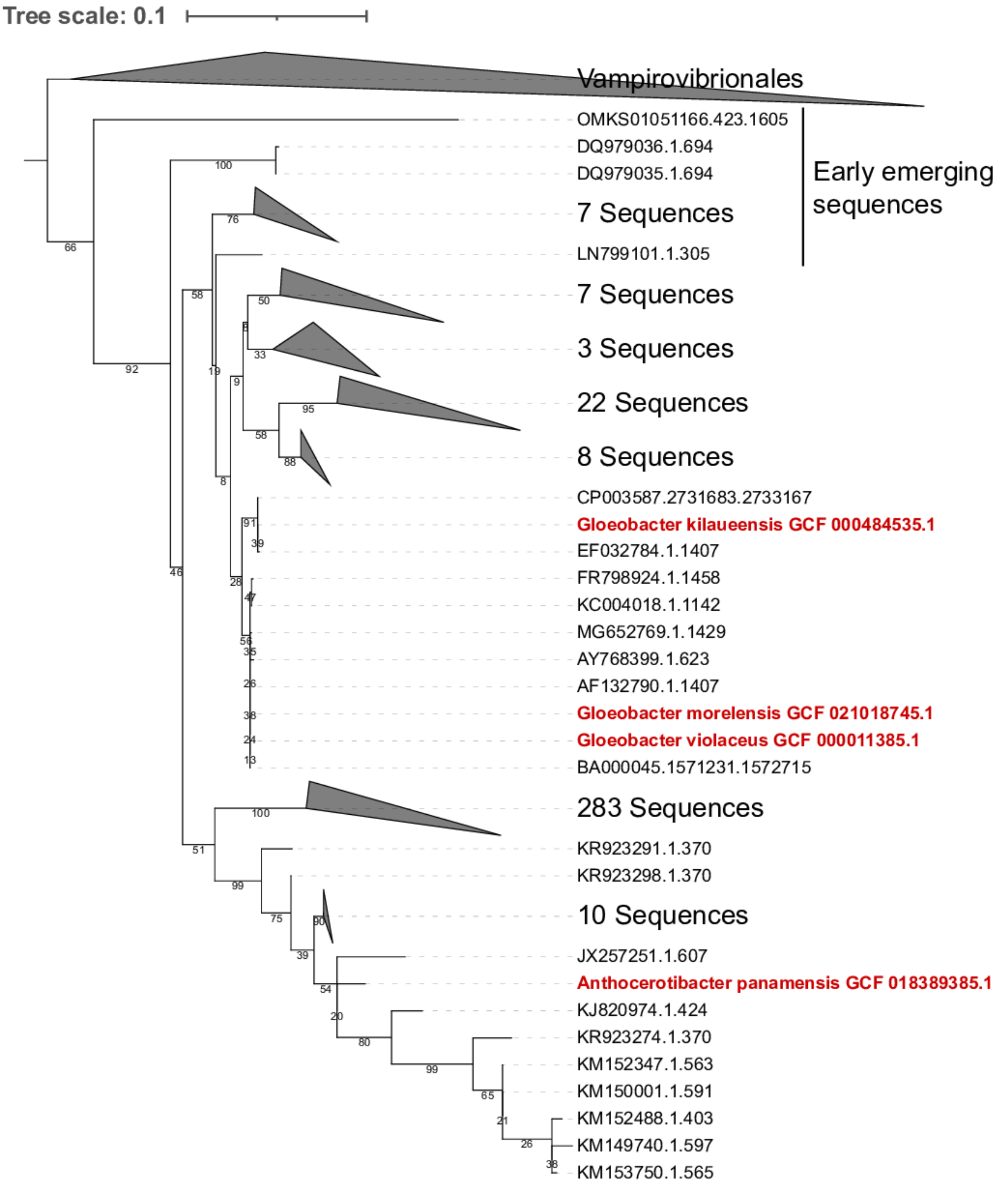


**Figure S5: Constrained SSU rRNA phylogeny. The reference phylogeny was built on ribosomal proteins with genomes from the NCBI, in red. This phylogeny constrains the SSU rRNA phylogeny, computed with RAxML [14] under the GTRGAMMA model.**

This SSU rRNA phylogeny shows a high diversity of unsequenced *Gloeobacterales* organisms. Eleven sequences are located at the base of the tree and might represent the earliest emerging Cyanobacteria known to date.

### 8. Average nucleotide identity comparisons

Average Nucleotide Identity (ANI) is used to define clusters of species, according to a 95% threshold.

**#ANI**
**#PATH: ANI/**
**$ nextflow run ANI.nf --genome=genome --list=list --cpu=20**
**#CPU time: 2m 13s**

The ANI matrix produced by fastANI indicates an important difference between *Gloeobacterales* genomes. The ANI values are all below 78 %, with the exception of the 92.8% between *Gloeobacter-morelensis*_MG652769 and *Gloeobacter-violaceus*_PCC7421.

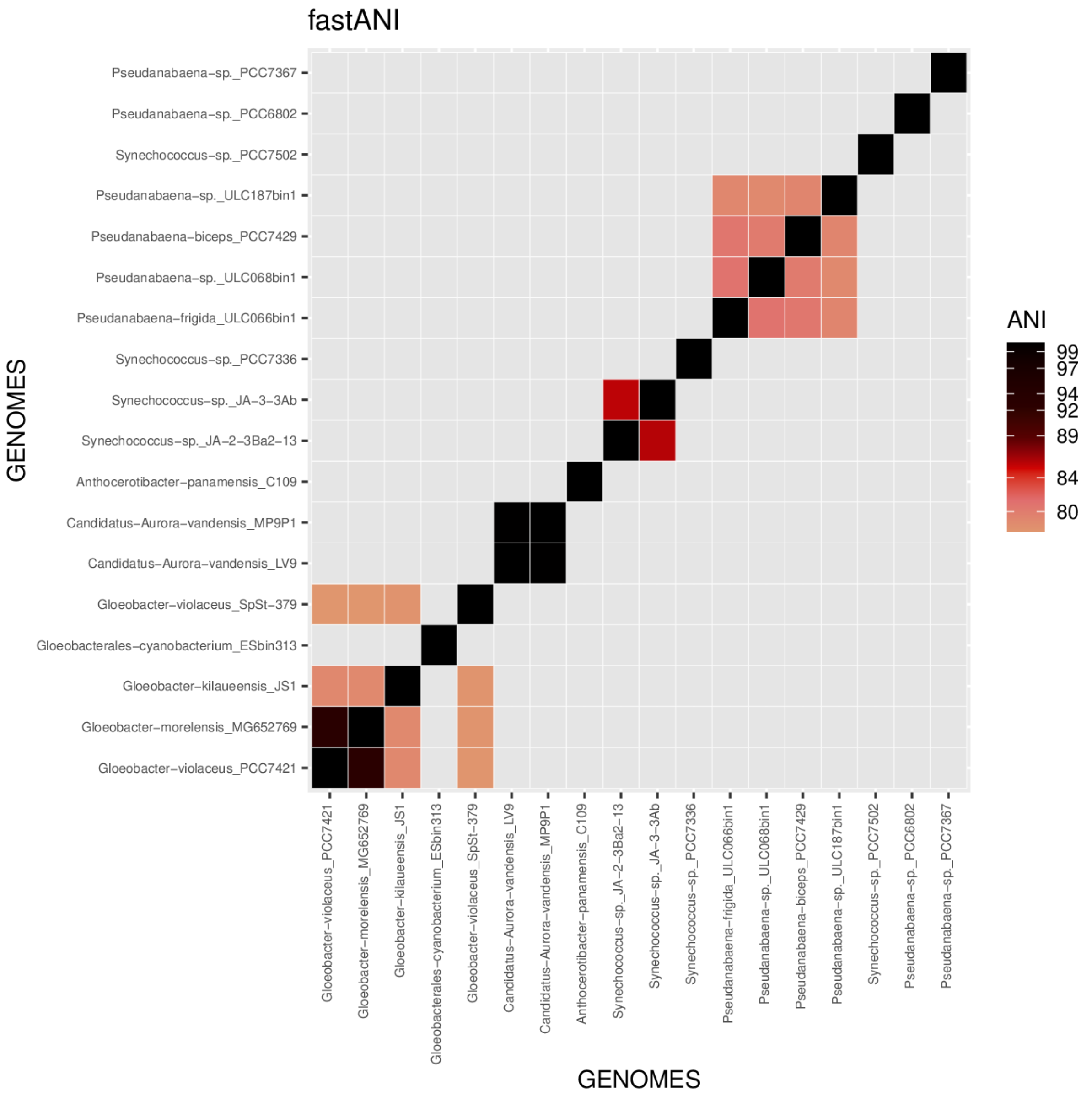

**Figure S6: Average nucleotide identity heatmap. Values below 78% are not plotted in the heatmap.**

### 9. Metabolic modeling

The metabolic modeling is used to map KEGG pathways on the genomes.

```
#Modelling
#PATH: /scratch/ulg/bioec/lcornet/GLOEO/metabo
$ nextflow run Metabolic.nf --infile=infile --mode=modelling --list=list \
--cpu=20 --kegg=/scratch/ulg/GENERA/Databases/ANVIO/kegg/
#CPU time: 1h 24m 59s
```

The workflow uses the anvi_metabolism script of Anvi'o [12].

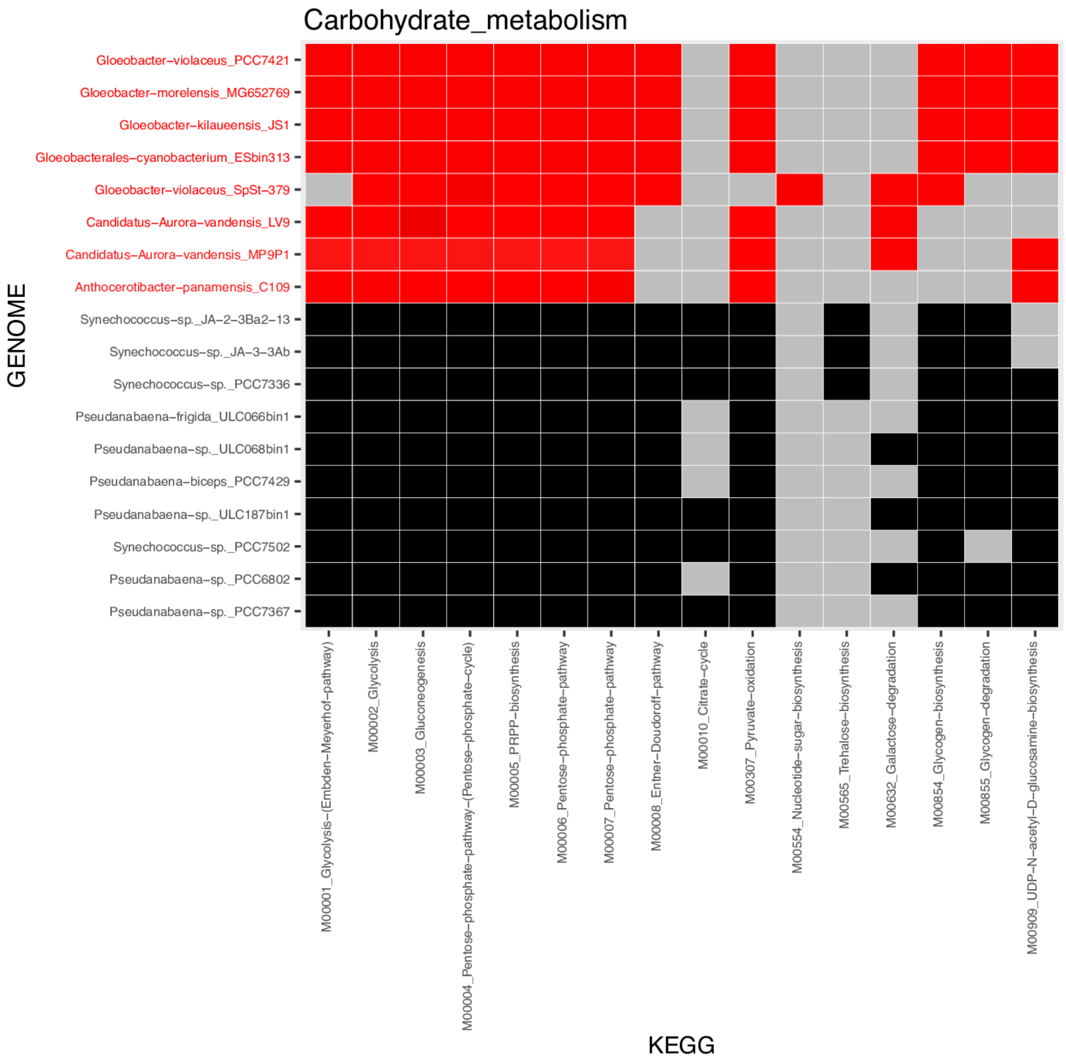

**Figure S7**: **Carbohydrate metabolism KEGG pathways. Black or red colour each indicates the presence of the pathways, according to Anvi'o [12] threshold; grey indicates the absence. *Gloeobacterales* genomes are indicated in red.**

Figure S7 shows an absence of the citrate cycle M000_10 (KEGG id), https://www.genome.jp/module/M00010, in all *Gloeobacterales* genomes. Beyond the carbohydrate metabolism, the workflow also produced plots for amino acid metabolism, biosynthesis of terpenoids and polyketides, energy metabolism, glycan metabolism, lipid metabolism, metabolism of cofactors and vitamins, and nucleotide metabolism.
